# Flexible and open-source programs for quantitative image analysis in microbial ecology

**DOI:** 10.1101/2022.09.23.509172

**Authors:** Alexis L. Pasulka, Jonathan F. Hood, Dana E. Michels, Mason D. Wright

## Abstract

Epifluorescence microscopy is an essential tool for obtaining reliable estimates of the abundance of marine microorganisms including viruses. However, computational analysis is required to gain consistent and quantitative data from digital microscopy images. Many imaging programs are proprietary and cost-prohibitive. The currently available free imaging programs are often platform specific and/or lack the flexibility to analyze microscopy images from natural samples, such as the planktonic environment, which can contain challenges such as debris and high background signals. Here we describe two MATLAB-based open-source image analysis programs that work across computer platforms and provide the tools to analyze a range of image types and cell sizes with a user-friendly interface. The Microbial Image Analysis (MiA) program aims to provide flexibility for the selection, identification, and quantification of cells that vary in size and fluorescence intensity within natural microbial communities. The Viral Image Analysis (ViA) program aims to provide an effective means for quantifying viral abundances from epifluorescence images as well as enumerating the intensity of a primary and secondary stain. In this paper, we provide an overview of the functionality of the MiA and ViA programs and highlight specific program features through several microbial image case studies.

## Introduction

Direct measurements of microbial abundance and biomass are critical for accurately characterizing the distribution of microorganisms (e.g., viruses, bacteria, phytoplankton, and microzooplankton) across marine ecosystems and their contributions to biogeochemical cycles in the ocean (Miloslavich et al. 2018, Khachikyan et al. 2019). Epifluorescence microscopy is a cornerstone of marine microbiology research (e.g., Hobbie et al. 1977, Weinbauer and Suttle 1997, Noble and Fuhrman 1998, Sherr and Sherr 1983) and has enabled scientists to visualize marine microbes across a wide range of sizes (e.g., <0.2 μm-200 μm). In addition to quantifying the abundance, biomass, and size structure of natural marine microbial communities (e.g., Patel et al. 2007, Christaki et al. 2011, Pasulka et al. 2013, Taylor et al. 2012, 2015), epifluorescence microscopy has been used to gain insight into particular taxonomic groups (via fluorescent *in situ* hybridization – FISH; Pernthaler and Amann 2004), growth rates (Hamasaki et al. 2004), microzooplankton grazing rates (Sherr et al. 1987), trophic modes (Caron 1983), and to determine active members of a microbial community (via substrate analog probing; Hatzenpichler et al. 2014, Samo et al. 2014). Automated quantitative imaging devices (e.g., Imaging FlowCytobot; Olson and Sosik 2007, Sosik and Olson 2007) are improving the spatiotemporal resolution over which marine microbial communities can be characterized and can help lead to an improved global plankton observation effort (see Lombard et al. 2019 for review of current technology). However, these efforts are not meant to replace precise, fine-scale, and high-quality local sampling conducted during oceanographic cruises or as part of site-specific observation sampling projects. In addition, super-resolution fluorescence microscopy approaches are changing our ability to visualize viruses and their interactions (Castelletto and Boretti 2021), but conventional wide-field fluorescence microscopes are still used to determine the abundance of viruses from environmental and culture samples (e.g., Turzynski et al. 2021 and sources within). Therefore, efforts are needed to continue integrating the visualization of microorganisms within discrete studies to gain comprehensive insight into how marine microbial communities are structured and their influence on marine ecosystem functions (Sebastian and Gasol 2019).

While microbial ecologists have used microscopy to visualize microbial communities for decades, advancements in microscope, camera, and computing technology have made digital image analysis a more common and essential tool (Wollman and Stuurman 2007, Waters 2009, Waters and Wittman 2014, Wait et al. 2020). Image analysis software programs exist, but many are proprietary and can be cost prohibitive (e.g., Imaris, ImagePro). Free programs such as ImageJ (imagej.nih.gov) and CellProfiler (McQuin et al. 2018, Carpenter et al. 2006) can be valuable for culture and larger-cell applications, but many lack the flexibility and customization needed to analyze complex environmental samples and small-particles like viruses. Programs like Daime (Daims et al. 2006) are more applicable to environmental samples, but are platform specific (e.g., Windows and Linnux). Furthermore, the quantification of viral particles remains a challenge across all platforms due to their small size (e.g., Shopov et al. 2000, Barrero-Canosa and Moraru 2018). A few MATLAB-based open-source programs have been developed to track the movement of viral particles (Jaqaman et al. 2008, Lee et al. 2016, Wang et al. 2018), but an easy-to-use software for quantifying viral particle abundance and fluorescence from cultured and environmental samples does not exist. Therefore, as tools such as phageFISH (Allers et al. 2013, Barrero-Canosa and Moraru 2018) and viral BONCAT (Pasulka et al., 2018) are applied in natural communities, open-source image analysis tools are still needed.

Here we describe two MATLAB-based, open-source programs for analyzing epifluorescence microscopy images of microbial communities. The programs can be run through MATLAB (on a Mac or PC) or can be downloaded as executable programs and run through the freely available MATLAB runtime environment. MATLAB has a breadth of functions useful for analyzing digital microscopy images, but these are inaccessible to users without a working knowledge of coding in MATLAB. The two programs presented here put the functions of MATLAB analyses in the hands of the users in an easy-to-use manner with no prior knowledge of code required. The Microbial Image Analysis (MiA) program aims to provide flexibility for the selection, identification, and quantification of cells that vary in size and fluorescence intensity (natural or probe-conferred) within natural microbial communities. Additionally, MiA has a cell-ID feature that enables the user to define and classify regions of interest (ROIs) real-time during image analysis. The Viral Image Analysis (ViA) program aims to provide an effective means for quantifying viral abundances from epifluorescence images as well as enumerating the intensity of a primary (e.g., SYBR Gold) and secondary stain (e.g., biorthogonal non-canonical amino acid tagging [BONCAT] or FISH). Both programs enable the user to export data in easy-to-use formats, facilitating downstream analysis. Below we provide an overview of the functionality of the MiA and ViA programs and highlight specific program features through several case studies. The case studies include microscopy images of:

1. a natural mixed phytoplankton community to demonstrate the flexibility of ROI selection and the functionality of the cell ID feature,
2. a mixed culture of the dinoflagellate grazer *Oxyrrhis marina* and phytoplankton *Dunaliella tertiolecta* to illustrate the separation of populations based the cell size and spectral properties collected by the program, and
3. *Emiliania huxleyi* viruses (EhV) to explore the quantification of viral abundance (via SYBR Gold staining) and the detection of a fluorescence signal from amino acid tagging.

### MiA and ViA Packages

#### Installation and Requirements

The MiA and ViA programs can run either as a script inside the MATLAB software or as an executable outside of the MATLAB software. Both the source-code for the script and the executable can be downloaded from a public GitHub repository (see Methods for details). In order to run the program via the source code in MATLAB, MATLAB R2020a or later must be installed. In order to run the executable program, the latest MATLAB Runtime Environment must be installed. Comprehensive online documentation for the programs can also be found on the GitHub public repository (see Data Availability section for details).

#### Package Structure and Overview of Modules

Overall, the MiA and ViA programs are constructed with a series of object-oriented packages and classes (Figure 1). The packages are named according to their functionality and include “bfmatlab”, “Constants”, “Events”, “Figure”, and “Interfaces”. The external package “bfmatlab”, is part of the Bio-Formats program developed by the Open Microscopy Environment (www.openmicroscopy.org) for opening Zeiss-formatted images (e.g., .czi files) with slight modifications to allow for visible status updates in the MiA and ViA graphical user interfaces (GUIs). The “Constants” package was designed to hold any desired program-wide constants. Currently, only graphical constants are held in that package, including x- and y-spacing values, figure position arrays, and small to large font sizes. The “Events” package was created to hold any custom events for the program. At the present stage, only a minimalist EventData wrapper subclass object is required to pass along single-action values as EventData. The “Figure” package holds all general items related to figure creation or figure manipulation classes, including a class that creates a completely blank figure, a customized question dialog box, a customized file selection panel, and a custom status update panel. These classes were designed to be modular, and can be leveraged to more efficiently create new “Figure” or “Interface” classes. Within “Figure”, there is a sub-package entitled “Functions” designed to hold any additional functionality capable of manipulating or modifying existing graphics. Currently, the only file within this sub-package is a modified version of an external MATLAB FileExchange program “dragzoom.m” that gives the user various abilities when dealing with one or multiple axes objects. In the Mac version of each program, this package also has a “MacFix” sub-package, specifically for the post-Catalina OS on Mac devices which interferes with MATLAB’s “uigetfile” ability to select separate file extension objects. Within this sub-package is a modified version of a MATLAB FileExchange “uigetfile_with_preview.m”, employing an older version of MATLAB file interface that does not have the same communication protocol problem.

**Figure 1.**
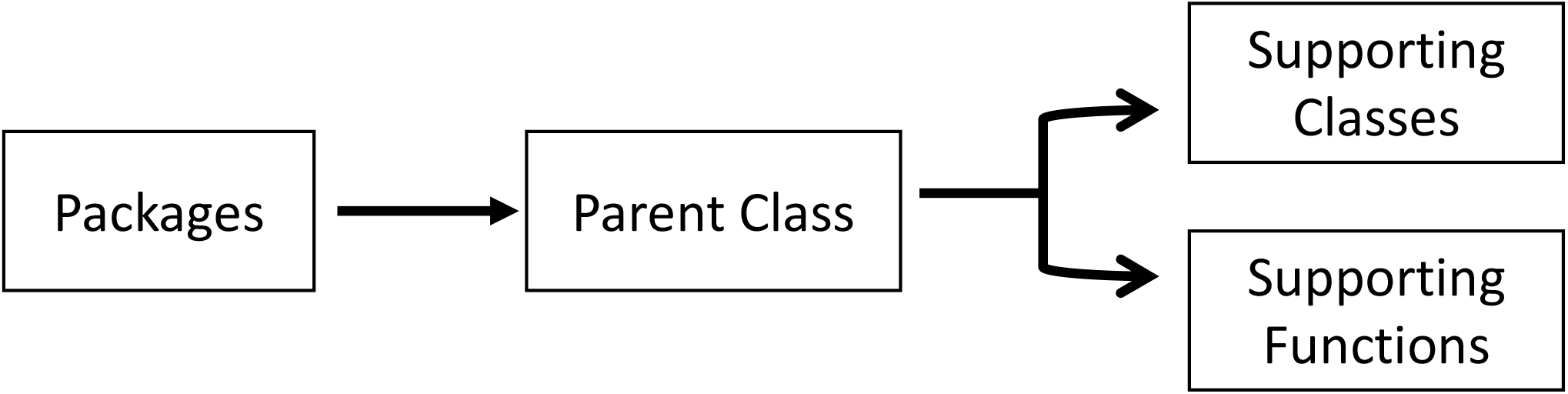
Overview of Program Structure.

The “Interfaces” package of both programs contains a series of classes. Each class is a subset interface (e.g., a full figure interface or an inset panel interface) designed to work with the primary interface “image_analysis.m” or “viral_analysis.m for MiA or ViA, respectively. Running the primary interface opens the full program. Within MiA, the classes include “analyze.m”, “bckgrnd_sub_interface.m”, “channel.m”, “manual_threshold_interface.m”, “roi_stats.m”, “select_channel.m”, and “roi_identification_interface.m”. Within ViA, the classes include “analyze.m”, “channel.m”, “manual_threshold_interface.m”, “roi_stats.m”, and “select_channel.m”. The “Interfaces” package varies the most between the MiA and ViA programs, and has minor differences between the PC and Mac versions. ViA has a sub-package “Functions” that holds functions necessary for the “Interfaces” to function. Currently, this sub-package contains a MATLAB FileExchange file by name of “findjob.m”, which extracts the underlying java object within a passed container or MATLAB GUI handle.

Collectively the structure describe above creates a simple user interface for both MiA and ViA. The MiA user interface displays the image in the middle of the panel, statistics on the left-hand side of the panel, and image options on the right-hand side of the panel (Figure 2). A series of dropdown menus provide the user the functionality to load images, select and modify ROIs, save data, and adjust display settings. After the user loads an image and assigns color channels, the program tools (Table 1) can be used in any order. Examples of how some of these tools can be used are described in case studies 1 and 2 below. The ViA program interface (Figure 3) differs from the MiA program interface because the processing of viral images requires less manual selection of ROIs and occurs in a specific order. The left-hand side of the interface displays the image (or images) and the right-hand side of the interface displays a series of panels that the user engages with in a sequential order to process a viral image (Table 2, Figure 3). Figure 3 is displaying the final processed viral image after subtracting the background, thresholding, and removing artifacts when either a single channel viral image is used (e.g., DNA signal; Figure 3A) or a dual-channel viral image is used (e.g., viral BONCAT signal; Figure 3B). More details for the steps are provided in Case Study 3 below.

**Figure 2.**
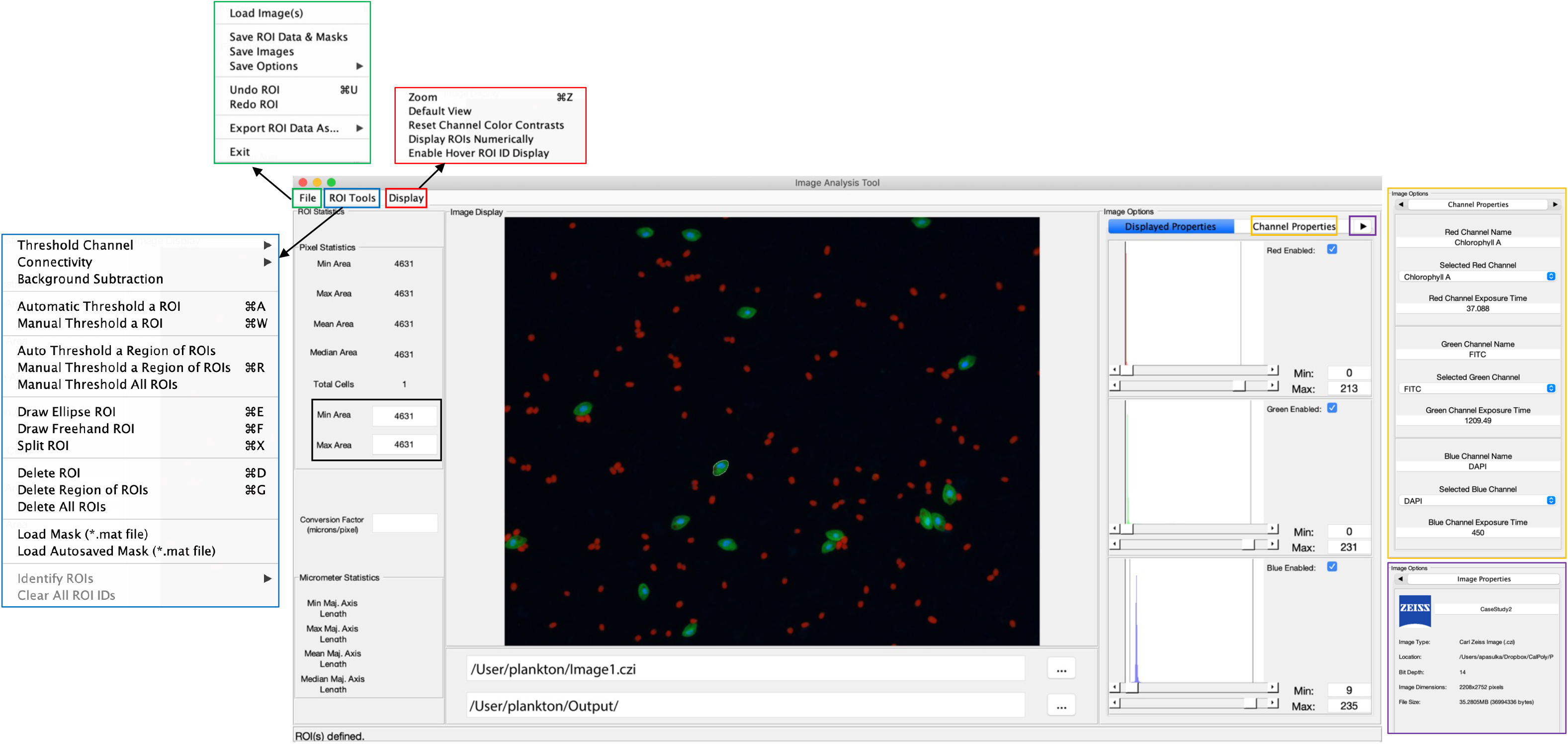
Overview of the MiA interface. The pop-outs on the left represent file options (green), ROI selection tools (blue), and display options (red). On the right, the image options are shown including display properties (inset), channel properties (yellow pop-out) and image properties (purple pop-out). The panel on the left also has a pixel size-selection feature (black square) that enables users to set a lower and upper pixel limit for ROIs. In the middle of the panel the image is visualized with a single cell outlined (e.g., region of interest; ROI). The file input directory and output directory are displayed just below the image and the statistics of the ROI (in pixels) are displayed just to the left of the image. If a conversion factor is added, the statistics will also be displayed in micrometers (μm).

**Table 1.**
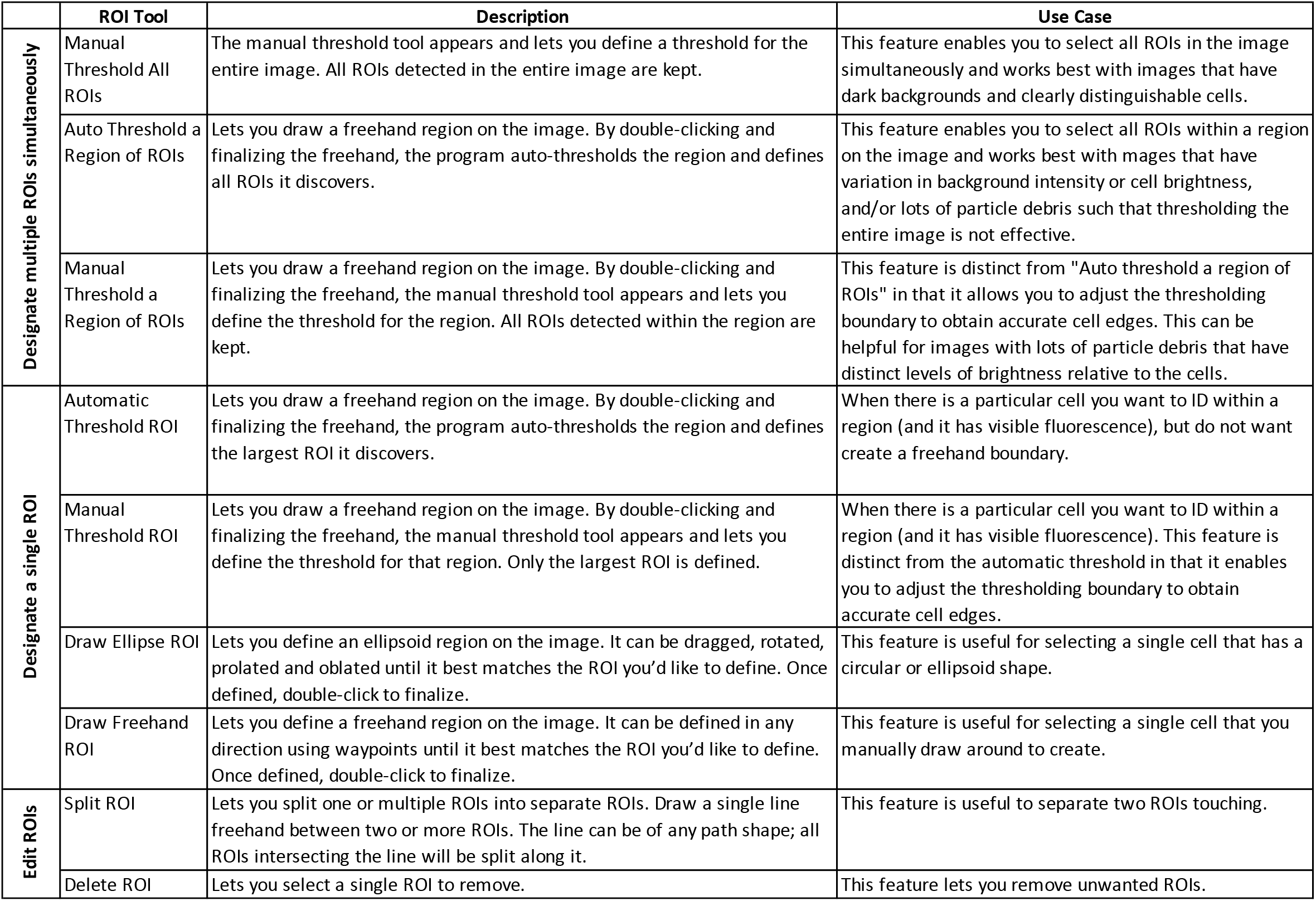
Description and use case for available ROI Tools for MiA

**Figure 3.**
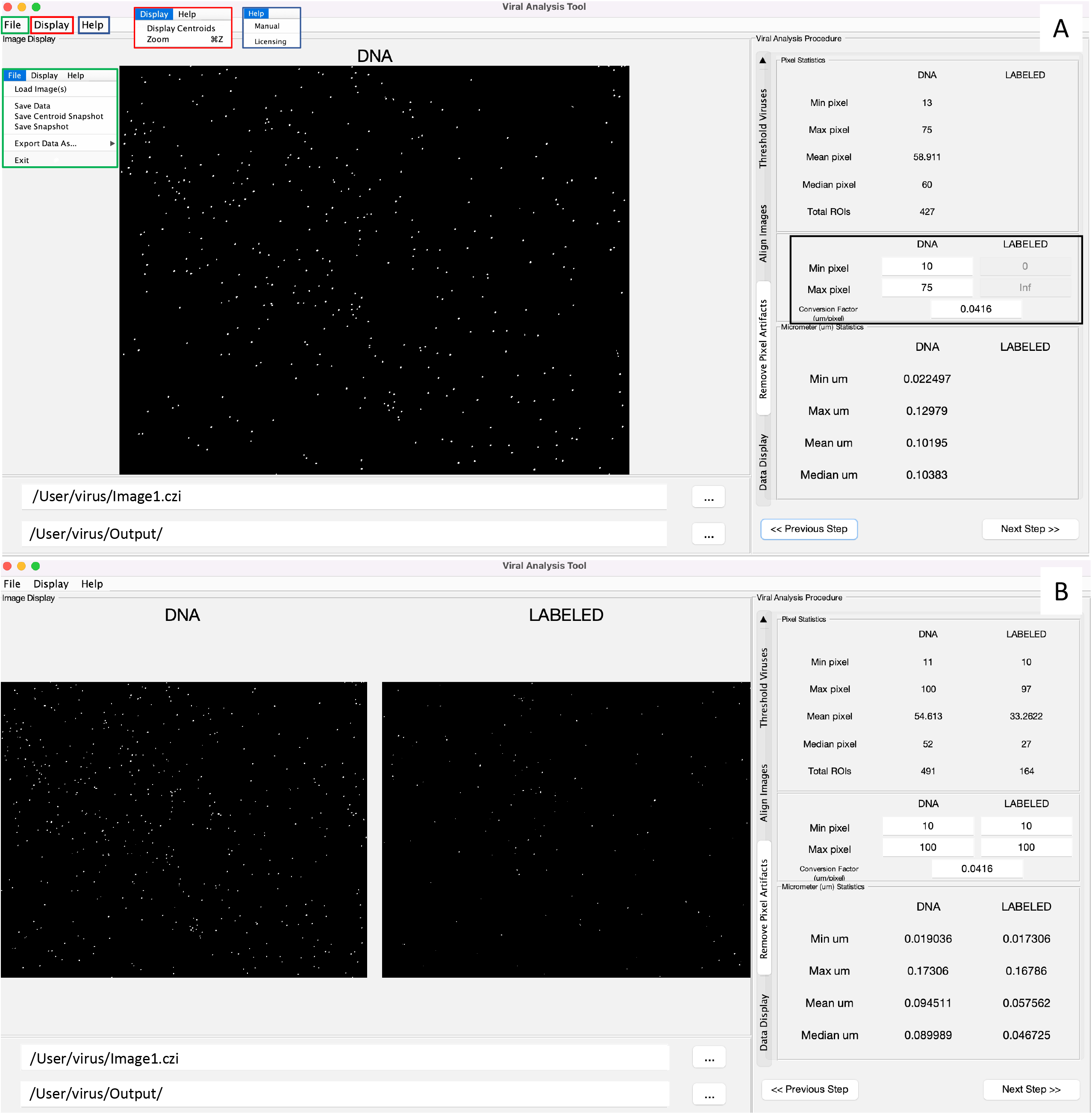
Overview of the ViA interface. The left hand side displays a single image if only one channel is loaded (A) or a double image if two channels are loaded (B). The right-hand side contains a series of panels that are used in a sequential order and details of the processing on each panel can be found in table 2. Shown here is the panel in which the user can decide the min and max pixel range of interest (black square), which then displays the statistics of the viral particles after the processing steps. The file input directory and output directory are displayed just below the image and the insets in panel A show the drop-down menus including the file menu (green), the display menu (red) and the help menu (blue).

**Table 2.**
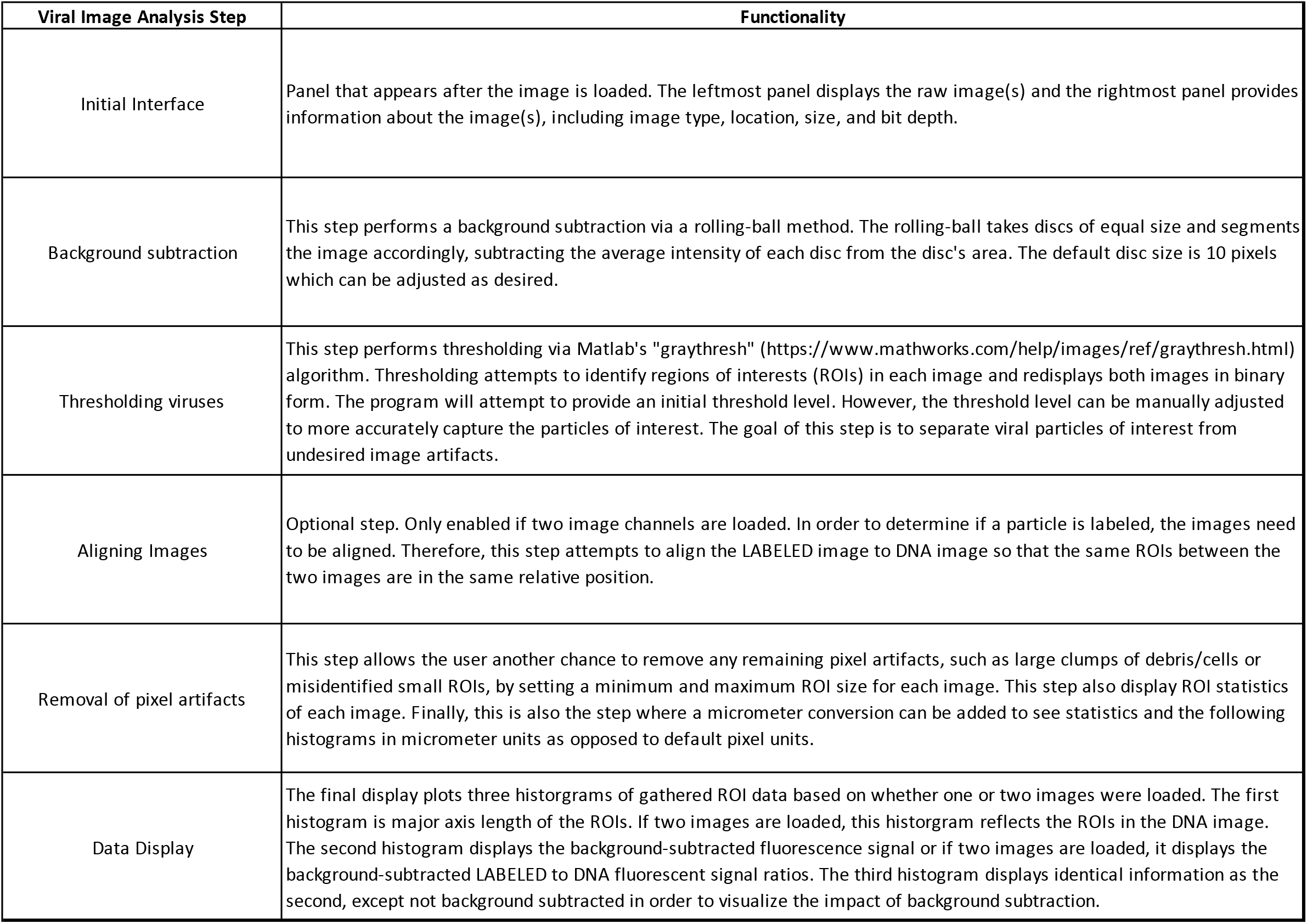
Description of steps required for processing images within ViA

| Viral Image Analysis Step | Functionality |
| --- | --- |
| Initial Interface | Panel that appears after the image is loaded. The leftmost panel displays the raw image(s) and the rightmost panel provides information about the image(s), including image type, location, size, and bit depth. |
| Background subtraction | This step performs a background subtraction via a rolling-ball method. The rolling-ball takes discs of equal size and segments the image accordingly, subtracting the average intensity of each disc from the disc's area. The default disc size is 10 pixels which can be adjusted as desired. |
| Thresholding viruses | This step performs thresholding via Matlab's "graythresh" (https://www.mathworks.com/help/images/ref/graythresh.html) algorithm. Thresholding attempts to identify regions of interests (ROIs) in each image and redisplay both images in binary form. The program will attempt to provide an initial threshold level. However, the threshold level can be manually adjusted to more accurately capture the particles of interest. The goal of this step is to separate viral particles of interest from undesired image artifacts. |
| Aligning Images | Optional step. Only enabled if two image channels are loaded. In order to determine if a particle is labeled, the images need to be aligned. Therefore, this step attempts to align the LABELED image to DNA image so that the same ROIs between the two images are in the same relative position. |
| Removal of pixel artifacts | This step allows the user another chance to remove any remaining pixel artifacts, such as large clumps of debris/cells or misidentified small ROIs, by setting a minimum and maximum ROI size for each image. This step also display ROI statistics of each image. Finally, this is also the step where a micrometer conversion can be added to see statistics and the following histograms in micrometer units as opposed to default pixel units. |
| Data Display | The final display plots three histograms of gathered ROI data based on whether one or two images were loaded. The first histogram is major axis length of the ROIs. If two images are loaded, this histogram reflects the ROIs in the DNA image. The second histogram displays the background-subtracted fluorescence signal or if two images are loaded, it displays the background-subtracted LABELED to DNA fluorescent signal ratios. The third histogram displays identical information as the second, except not background subtracted in order to visualize the impact of background subtraction. |

### Image Analysis Examples and Workflow

In this section, we have selected a range of epifluorescence images to showcase the capabilities of these imaging programs including 1) flexible options for region of interest (ROI) selection, 2) the ROI identification tool, 3) an example of the quantitative data that gets extracted from ROIs, and 4) several examples of how this data can be used to characterize microbial community structure (via abundance or size) or quantify a fluorescence signal (e.g., fluorescence in situ hybridization signal). Materials for each case studying including images and data generated from the images are all available on the public GitHub repository (see Data Availability section for details).

### Case Study 1 – Natural Plankton Community

#### Flexible options for region of interest (ROI) selection

The MiA Program has a variety of ROI selection options (Table 1) that enable the user to accurately and efficiently select cells across a range of image types. Analyzing images produced from complex environmental samples can be challenging due to varying degrees of fluorescence signals across cells and background signal from debris (Figure 4A). Therefore, these types of images often require different analysis strategies than images with consistent cell types and dark backgrounds (e.g., Case Study 2). The program allows for seamless toggling between different cell-selection approaches to best meet the needs of each area of an image. The user can also adjust contrast within any channel real-time during analysis (Figure 4B), which does not alter the data in any way, but gives users the ability to intensify the signal of a dim cell for the purposes of cell selection.

**Figure 4.**
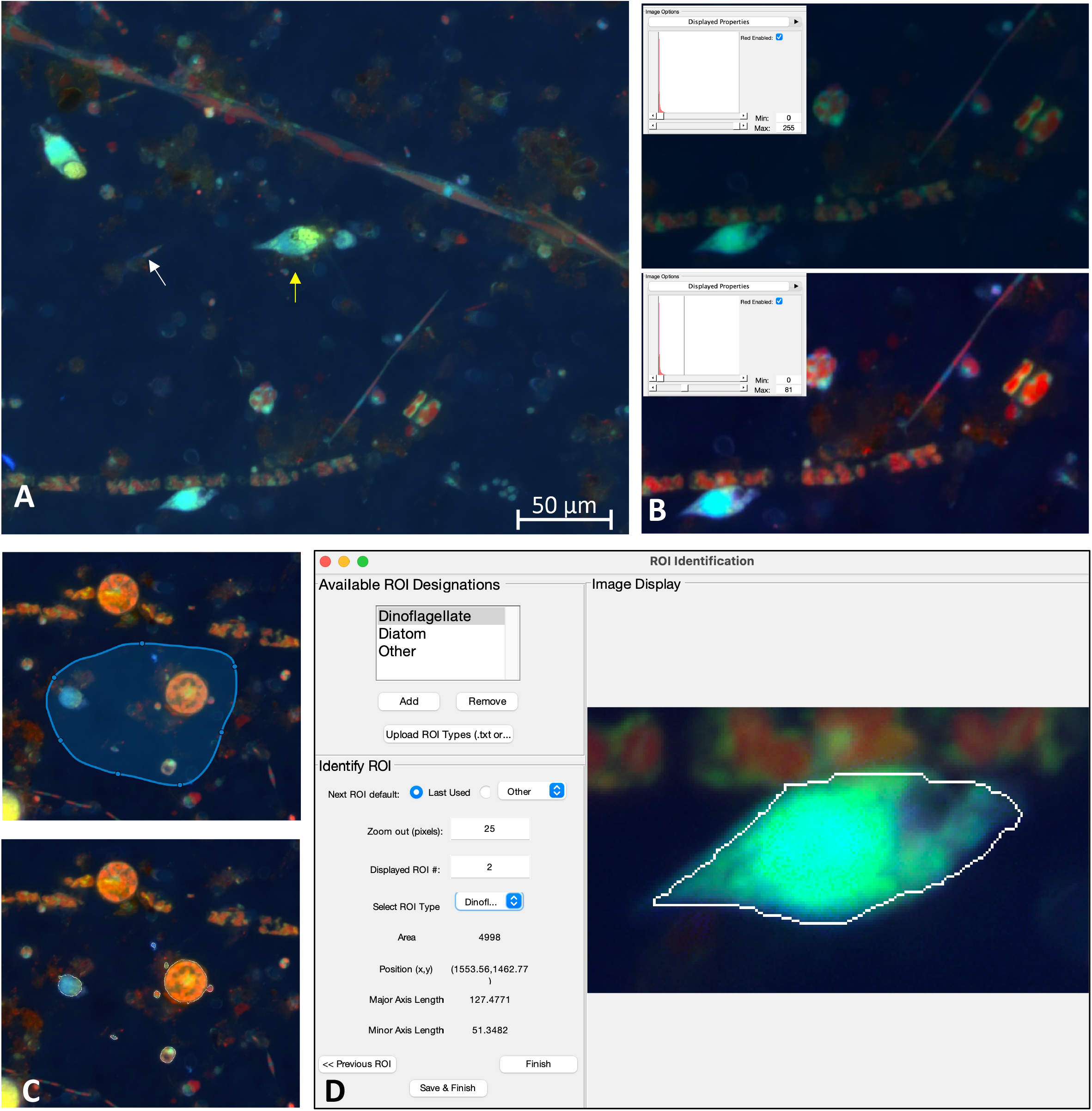
A) Example image of natural plankton community with fainter diatom cells (white arrow) and brighter dinoflagellate cells (yellow arrow). B) An example of contrast adjustments available for real-time cell selection showing how altering the red signal (inset) enhances the ability to see cells for ROI selection (bottom image displays enhanced signal). C) Example of regional thresholding – both the selected region (top) and outlined cells within the region (bottom). D) ROI Identification window with a dinoflagellate outlined.

While MiA enables thresholding cells across the entire image at once, uneven backgrounds can make whole-image approaches problematic. Therefore, regional thresholding is particularly valuable for environmental images (Figure 4C). To add additional flexibility, the user can select the channel to be used by the thresholding algorithm for defining ROIs. In addition, the program offers users the ability to select individual cells (e.g., single ROI selection) or carry out free-hand drawing. If two cells are close together and are incorrectly selected as one cell, the ‘split cell’ feature enables the user to easily separate the cells (see Case Study 2 and Figure 5B for details). The program also offers a number of ROI removal options. Users have the option to delete a single ROI, multiple ROIs within a selected region, or all ROIs. In additional to ROI removal, there is a pixel size-selection feature that enables users the ability to set a limit and remove small cells (or even image artifacts) or set an upper limit and remove large cells (Figure 2). The background subtraction feature, with several different strategies to choose from, can be used for more complicated images of natural microbial communities. It is important to note that because background subtraction has the potential to alter the data, the original and background subtracted data are provided upon data export. While users can only visualize three channels at a time during image analysis (e.g., Red, Green, Blue), if additional channels exist, users can switch between the channels that are visualized during image analysis and ROI selection. Furthermore, data from all channels (not just those visualized) can be exported at the end of an image analysis session using the mask file (see ‘Saving Options’ for details).

**Figure 5.**
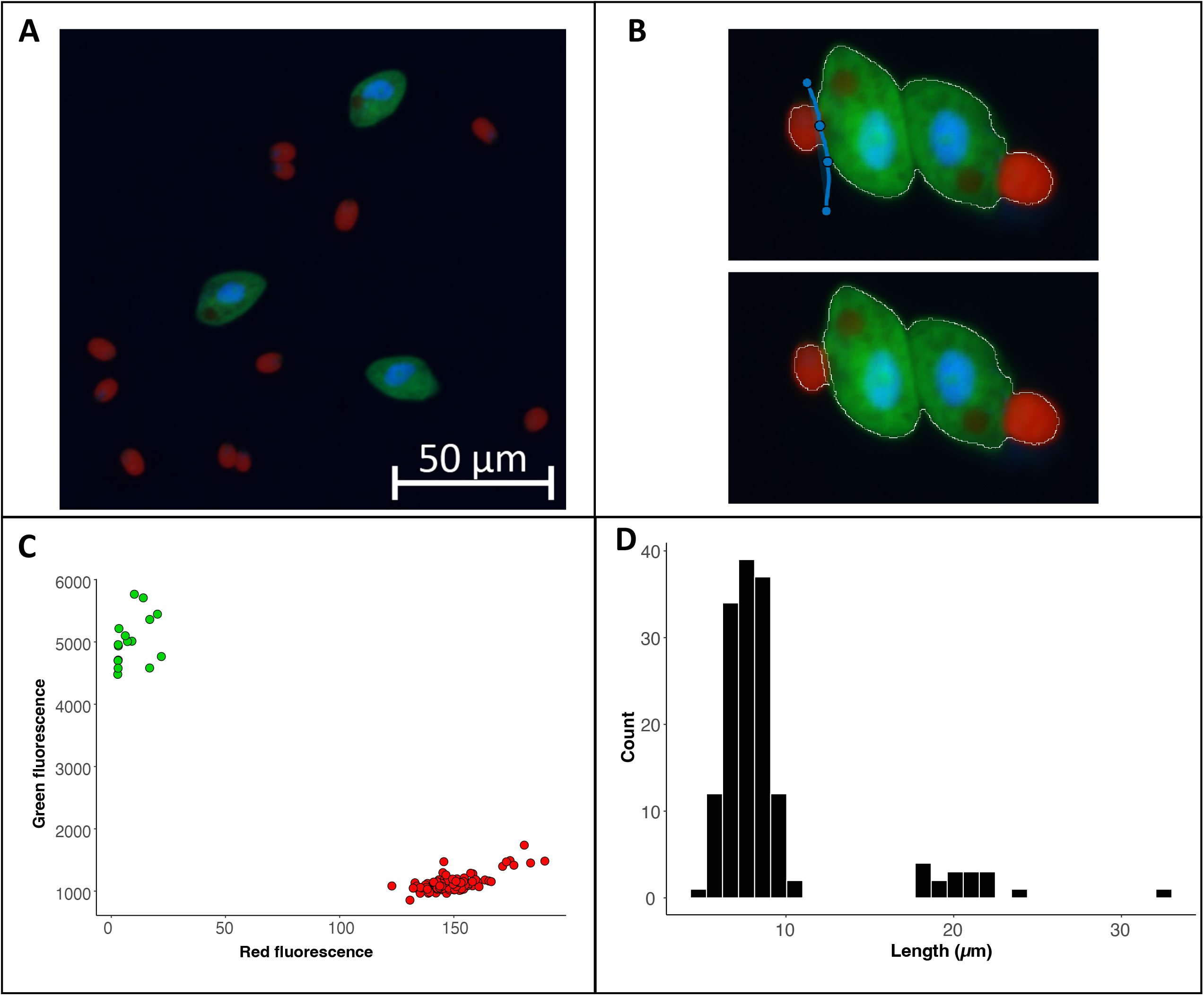
Example image of from a culture of *O. marina* (green cells) fed *D. tertiolecta* (red cells) (A). The split cell feature can be used to separate cells that were outlined as a single cell (B). The data exported from the program can be used to visualize the fluorescence or size data collected within each region of interest. For this case study we have used R to visualize the separation of cells by fluorescent signal with the *D. tertiolecta* (red circles) having a higher red fluorescence signal and the O. marina (green circles) having a higher green fluorescent signal (C). In addition, we have visualized the size distribution of cells from the image with the smaller cells representing the prey *D. teriolecta* and the larger cells representing the dinoflagellate grazer *O. marina* (D).

#### ROI identification feature (ROI ID)

While cell fluorescence and/or cell size can be used to separate populations of interest using the exported data after image analysis (see Case Study 2), specific types of cells from mixed, complex communities can be more challenging to identify from these types of data signals. Therefore, while carrying out image analysis the user has the option to manually identify and classify cells (Figure 4D). The user can enter different names by which they would like to identify cells (e.g., dinoflagellate and diatom). The feature enables users to identify only one type of cell (e.g., diatoms), or identify multiple cell types. While programs have been developed to obtain automated taxonomic classification and quantitative data from epifluorescence images (e.g., Hense et al. 2008, Schulze et al. 2013, Colin et al. 2017), these programs require large reference training sets. This scale of image analysis is not always required, nor feasible; therefore, there is still a need for manual image analysis for smaller-scale studies.

#### Saving options

Working with microscopy images from a natural environment can be time-consuming and require multiple iterations. Therefore, the MiA program enables users to save a ‘mask’ file (.mat), which is a small file containing information about the masks or regions of interest identified by the user. Masks can be easily loaded and modified during any analysis session. The program also has an autosave feature that saves a mask file in the event that there is a computer issue during analysis. When image analysis is complete, the user can export the data from their regions of interest. For each ROI, the data includes the ROI number, ROI identification (if designated), the fluorescence intensity (min, mean, and max) of the ROI in each color channel, the area, length, width and perimeter (all in pixels) of the ROI, and the x-y coordinates of the ROI on the image. If background subtraction was used (an option tool in the ROI Tools menu; Figure 2), the data also includes background subtracted fluorescence values, in addition to the original data. A ‘Data Summary’ sheet also gets saved as a second sheet in the file. This sheet contains the data visible in the ‘ROI Statistics’ panel of the primary program interface including total cells and min/max/mean/median ROI area in pixels. In addition, if a pixel to micron conversion factor was included, these statistics are also displayed in microns. While carrying out image analysis, the user can also save snapshots of the image. The snapshots maintain the current contrast adjustments and can be saved with or without outlines around the identified cells.

### Case Study 2 – Culture of phytoplankton (*Dunaliella tertiolecta*) and grazer (*Oxyrrhis marina*)

#### Flexible options for region of interest (ROI) selection

Images collected from plankton cultures, which typically have dark and even backgrounds, provide an opportunity to demonstrate a straightforward ROI selection process (Figure 5A). Additionally, the fluorescence data collected from these ‘clean’ digital images provides an opportunity to demonstrate how to separate populations and glean information from the ROI data post image analysis. Global thresholding can be used to threshold cells across the entire image. Using this feature, cells in close proximity to one another often get selected as a single ROI. In these cases, the split cell feature enables users to quickly and accurately separate individual cells by simply drawing a line through the ROI along the cell border (Figure 5B).

#### Quantification of populations based on size or fluorescence signal

While manual ROI identification, as demonstrated in Case Study 1, can be valuable for complex images, differentiating cell types by size or fluorescence signal can enable higher throughput means of ROI identification when working with images that have dark backgrounds and clear cell borders. Users can easily work with the exported data as part of the image analysis program. For Case Study 2, the phytoplankton and grazer image, we can separate the populations based on the red to green signal (Figure 5C) because the phytoplankton cells have a chlorophyll signal that the heterotrophic grazers do not. However, the signal used to separate populations can also come from an artificial label through FISH or BONCAT (e.g., Michels et al. 2021). The exported data also enables users to explore the size structure of the microbial community (Figure 5D) and quantify the concentration of different cell types (example R code is available with the case study on the GitHub repository).

### Case Study 3 – Viral Image Analysis

#### Quantifying the abundance of viral particles

Digital image analysis has been shown to be more efficient and accurate compared to microscopy-based estimates for enumerating viral particles from environmental samples, (Shopov et al. 2000, Chen et al., 2001, Barrero-Canosa and Moraru 2018). ViA, built from the pipeline developed by Pasulka et al. (2018), is distinct from MiA because the ROI selection is designed to deal with the challenges of imaging small particles. Therefore, images are processed in a sequential manner (Table 2), but the user can modify the settings for each step.Single-channel images (Figure 3A) or dual-channel images (Figure 3B) can be loaded and processed. Dual channel images (discussed in more detail below) may be of interest when quantifying a FISH or BONCAT signal within viral particles. When images are loaded, the images can be assigned to either the “DNA” or the “LABELED” signal (Figure 3). It is important to note that the image assigned to the “DNA” signal is considered the true signal and is what will be used to define viral particles. The image processing steps are outlined in Table 2 and discussed in more detail below.

In digital images, the background adds to the signal of interest (Waters and Wittman 2014). Since viral particles can vary in their fluorescence signal intensity (particularly when imaging a natural viral community), the background must be measured and subtracted from the intensity values of the pixels containing the signal of interest. Therefore, background subtraction is a critical first step in image processing using the ViA program (Table 2, Figure 6). The program uses a rolling-ball subtraction method. In short, a background value is determined for every disc (default disc size = 10 pixels), and the average intensity of each disc is subtracted from the disc’s area. Therefore, spatial variations in background intensity are easily accounted for in this approach and do not influence the ability to detect viral particles across the image.

**Figure 6.**
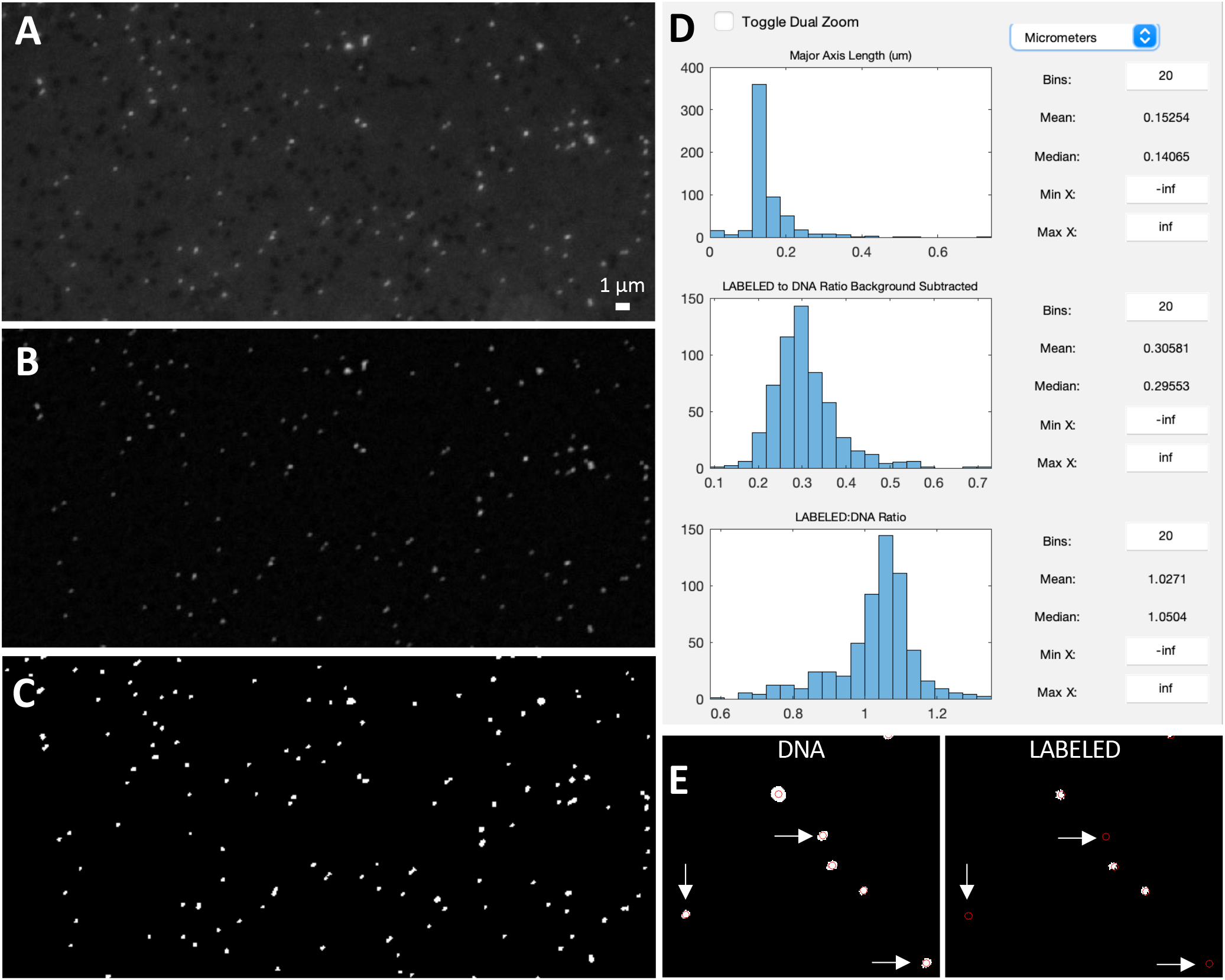
Example image of viruses before background subtraction (A), after background subtraction (B), and after thresholding (C). Data display panel (D) showing the size and fluorescent ratios of viral particles. Zoomed in regions of a ‘DNA’ and ‘LABELED’ images (E) showing centroids around viral particles (as defined by the DNA image). White arrows indicate viral particles that do not have a LABELED signal.

Thresholding is then used to identify the viral particles in the image (Table 2, Figure 6). In this type of image segmentation, the image is converted from a grayscale image (Figure 6B) to a binary image (e.g., black and white; Figure 6C). In this way, thresholding is used as a way to select ROIs (i.e., white regions) while ignoring the rest of the image (i.e., black regions). The program provides an initial threshold level using Otsu’s method (Otsu 1979), but the user has the flexibility to adjust this level to find a balance between image artifacts and viral particles.

In the final step of image processing, small and/or large image artifacts can be removed (Figure 3). By selecting a minimum and maximum pixel size, the user can alter the number of particles that are considered real. When these values are adjusted, the data displayed in the statistics panel will also change (Table 2, Figure 3). At this point of image processing, ‘Total ROIs’ should reflect viral particle abundance for the image. If the user inputs a micrometer (μm) to pixel conversion (also available at this step), then the size statistics (min, max, median, and mean) of those particles in μm is also provided. While the user can record this information from the panel, these statistics are exported in the “summary sheet” upon data export.

Before exporting the data, the final data display panel may be useful for visualizing the size distribution of viral particles (Table 2, Figure 6D). This image processing pipeline has been used to visualize and quantify viral particles ranging from 50-200 nm using SYBR Gold (Pasulka et al. 2018). While the ability to resolve two individual particles from one another is set by the objective, the pixel resolution is set by the CCD camera; therefore, careful consideration of camera capabilities is critical for downstream analyses of viral particles. However, it is important to keep in mind that the size of the fluorescent signal is not the actual size of the viral particles (see Figure S5 in Pasulka et al. 2018). The final data display also provides information about the intensity of the fluorescence signal within the particles (Figure 6D), but this data may be more useful if two images are loaded (see below for more details). It is important to note that while the panels are meant to be used in a sequential order the first time an image is processed, the user can go back to any panel and adjust settings as needed.

#### Detecting a fluorescent signal in viral particles

Approaches such as phageFISH (Allers et al. 2013) and viral-BONCAT (Pasulka et al. 2018) provide the ability to quantify the abundance of particular types of viruses and/or monitor viral infection dynamics, respectively. However, digital image analysis is still needed in order accurately quantify the co-localization of fluorescence signals. The general processing of viral images is the same if one or two channels are loaded. However, when two image channels are loaded, the user can input different background subtraction values, threshold levels, and artifact removal settings for each image.

One additional step that gets activated when two image channels are loaded is the image alignment step (Table 2). Proper microscope alignment is critical for optimal image analysis. While nanometer differences between filter cube alignment do not pose a problem for larger cells (e.g., >1μm), these shifts can be problematic for sub-micron particles such as viruses, especially when you are interested in co-locating a fluorescence signal. Therefore, the alignment step is meant to ensure the images are properly aligned.

The artifacts removal panel will now show the particle statistics for both image channels (Figure 3B), which can be useful for determining how many particles are labeled. When working with two image channels, the channel labeled DNA is considered the image with the ‘real’ viral particles. To visualize how labeled viral particles match up with these DNA viral particles, the display centroids feature can be used (Figure 2, Figure 6E). This places red circles around all DNA-image defined ROIs on both images. The final data display panel is also a useful place to visualize this information, as the histograms now show the LABELED to DNA fluorescence ratio (Figure 6D). While the user is recommended to export the data and process the signal for labeled viruses according to other methods (e.g., Pasulka et al. 2018), the red to green fluorescence ratio (i.e., the DNA to LABELED ratio) was distinct in viral particles produced from a host grown in the presence of HPG relative to viral particles produced by an unlabeled host control culture. Therefore, the ratio of fluorescence signals can provide the user a quick peek of the level of labeling in a treatment if compared to an unlabeled control (Pasulka et al. 2018). The display shows both the raw fluorescence data and the background subtracted fluorescence data so the user can quickly visualize the effect of background subtraction on the signal (Figure 6D). Therefore, if changes are needed in the background subtraction step (or any step), they can occur during image processing. Furthermore, the histograms can be saved as an image for quick reference later.

## Discussion

MiA and ViA were designed as open-source microscopy image analysis programs (GNU General Public License version 3) that work on both PCs and MACs, are easy to use, and provide the tools to analyze a range of image types and cell sizes. While MiA and ViA are MATLAB-based programs, the user does not need to have any coding knowledge to use the programs. Furthermore, the executable versions are available for users who do not have access to the proprietary MATLAB software. Open-source imaging programs such as these are meant to provide transparency and reproducibility for data collection from microscopy images. Furthermore, the code is open-source, which encourages improvements as well as the flexibility for the community to take the program in new directions. While the field of marine microbiology is moving towards more automated image analysis (Benfield et al. 2007, Schulze et al. 2013, Colin et al. 2017), the ability to gain quantitative information from microscopy images with flexible ROI-selection options without needing to purchase expensive software and/or to develop large training sets is still needed. While the case studies presented here focus on marine microbial communities, the functionality of the MiA and ViA programs is broadly applicable to any field of microbial ecology for analyzing microscopy images from cultured or environmental samples.

### Limitations and Potential Developments

While the focus for the development of the first iterations of MiA and ViA was manual image analysis from complex environmental samples, some image analyses would benefit from a more routine and faster procedure. Therefore, future iterations of the program could run an analysis on batches of similar images after the user sets certain parameters (e.g., thresholding channel, thresholding level, etc.).

Program memory influences the size of the program and the speed at the which the program can be used. MiA and ViA currently have temporary memory during an imaging session. For example, loading new images during the same session retains some preferences, such as the directory of the last image selected and the order of set color channels for grayscale and CZI images. However, the programs do not currently maintain any settings between program instances. In future iterations of the program, the temporary memory could be used as a foundation to develop greater program memory and enable the user to save desired settings between sessions.

The data collected by the program is currently provided in an easy-to-use format, which enables the user flexibility with the types of downstream analyses they can perform in their program of choice (e.g., excel, R, python). However, the programs cannot currently be used to perform any statistics on the image data. Based on user needs, future iterations of the program could leverage MATLAB’s Statistics and Machine Learning Toolbox.

## Data Availability Statements

The programs, datasets, and code can be found in the following public GitHub repositories: https://github.com/PECO-CP/MiA and https://github.com/PECO-CP/ViA.

## Author Contributions

ALP conceptualized both programs, wrote the initial code used in ViA, was involved in program testing and improvements, and wrote the manuscript. MDW wrote the code for an early iteration of MiA and contributed to program improvements. JFH was the primary contributor to the code for MiA and ViA, played an active role program development and testing, and was involved in manuscript and manual writing. DEM contributed to the testing of MiA and ViA, contributed to manuscript and manual writing, and was involved in case study development. All authors revised and approved the manuscript.

## Methods

### Software

The open-source software described above is available online at https://github.com/PECO-CP/MiA and https://github.com/PECO-CP/ViA. Materials for all three case studies as well as detailed manuals are also available on this public repository.

### Sample preparation and fixation

For the natural plankton image, a surface water samples (75 mL) was collected at the Cal Poly Pier in Avila Beach, CA (35.1698° N, 120.7408° W) and preserved with alkaline Lugol’s solution (0.05% final concentration) followed by paraformaldehyde (PFA; 2% final concentration) and sodium thiosulfate (0.003% final concentration) using a modified protocol from Sherr and Sherr (1993). The preserved sample was fixed at 4°C for 24 hours prior to filtration. The sample was stained with proflavine (0.33% final concentration) and DAPI (0.05 μg/mL final concentration) prior to filtration. Samples were then filtered onto 8.0μm black polycarbonate filters, mounted onto glass slides with VectaShield mounting medium (Vector Labs) and kept frozen at −80°C until imaging. For the culture image, *Duniella tertiolecta* was added to a culture of *Oxyrrhis marina* as prey and minutes later the mixed culture was fixed with glutaraldehyde (0.5% final concentration) at 4°C for 24 hours. 10 ml of sample was filtered onto a 0.8 μm black polycarbonate filter, mounted onto glass slides with DAPI VectaShield mounting medium (Vector Labs) and kept frozen at −80°C until imaging. The virus image was prepped as in Pasulka et al. (2018). EhV207 (MOI of 5) was added to a culture of *E. huxleyi* (CCMP strain 374) in exponential phase. Upon host lysis, the sample was filtered through a 0.45 μm filter, fixed with glutaraldehyde (0.5% final concentration) for 15 min at 4°C, flash frozen in liquid nitrogen, and stored at −80°C. The sample was then spotted directly onto a Teflon printed glass slide (Electron Microscopy Sciences, PTFE Printed Slides) and air-dried. The sample was counterstained for 15 min with SYBR Gold (0.25% final concentration), washed with 0.02-μm filtered water, and air-dried prior to image analysis.

### Microscopy

Samples were analyzed with a Zeiss Axio Observer Z1 inverted epifluorescence microscope using a 20X (natural phytoplankton community and culture image) or 100X objective (virus image) using Zen Microscope Software. Digital images were acquired with a 6-megapixel CCD camera (Zeiss Axiocam 506 mono). The peak channel excitation and emissions wavelength/bandpass in nm were 365 and 445/50 for blue (DAPI-stained cells), 470/70 and 525/50 for green (fluorescence signal of proflavine and SYBR gold as well as autofluorescence signal of glutaraldehyde), and 440/40 and 675/50 for red (chlorophyll autofluorescence). For the natural phytoplankton community sample, 10 z-plane images were acquired for each fluorescence channel. The resulting z-stack images were subsequently combined using an extended depth of focus (EDF) algorithm within the Zen software (ZEN Blue 2.3) to create an in-focus image.

## Acknowledgements

We acknowledge Cal Poly’s Research, Scholarly, and Creative Activities Grant Program for supporting the development of the image programs and DEM as well as the Bill and Linda Frost Fund for supporting summer research experiences for both MDW and JFH. We would also like to thank all of the students who used and tested the program, including several supported by the Santa Rosa Creek Foundation. We would also like to acknowledge Polerecky et al. (2012) who’s open-source nanoSIMS data processing tool provided inspiration for this Matlab-based imaging program.

## References

Allers, E., Moraru, C., Duhaime, M. B., Beneze, E., Solonenko, N., Barrero-Canosa, J., Amann, R., Sullivan, M. B. (2013). Single-cell and population level viral infection dynamics revealed by phageFISH, a method to visualize intracellular and free viruses. Environ. Microbiol., 15: 2306–2318.

Barrero-Canosa, J. and Moraru, C. (2018). PhageFISH for monitoring phage infections at single cell level. In Martha R. J. Clokie et al. (eds.), Bacteriophages: Methods and Protocols, Volume IV, Methods in Molecular Biology, vol. 1898.

Benfield, M., Grosjean, P., Culverhouse, P., et al. (2007). RAPID: research on automated plankton identification. Oceanography, 20: 172–187.

Carpenter, A.E., Jones, T.R., Lamprecht, M.R., Clarke, C., Kang, I.H., Friman, O., Guertin, D.A., Chang, J.H., Lindquist, R.A., Moffat, J., Golland, P., Sabatini, D.M. (2006). CellProfiler: image analysis software for identifying and quantifying cell phenotypes. Genome Biology, 7: 17076895

Caron, D. A. (1983). Technique for the enumeration of heterotrophic and phototrophic nanoplankton, using epi-fluorescence microscopy, and comparison with other procedures. Appl. Environ. Microbiol., 46: 491–498.

Castelletto, S. and Boretti, A. (2021). Viral particle imaging by super-resolution fluorescence microscopy. Chemical Physics Impact., 2: 100013.

Chen, F., Lu, J-R, Binder, B.J., Liu, Y-C., Hodson, R.E. (2001). Application of digital image analysis and flow cytometry to enumerate marine viruses stained with SYBR Gold. Appl. Environ. Microbiol., 67: 539–545.

Christaki, U., Courties, C., Massana, R., Catala, P., Lebaron, P., Gasol, J.M., Zubkov, M.V. (2011). Optimized routine flow cytometric enumeration of heterotrophic flagellates using SYBR Green I. Limnol. Oceanogr. Methods, 9: 329–339.

Colin, S., Coelho, L. P., Sunagawa, S., Bowler, C., Karsenti, E., Bork, P., Pepperkok, R., de Vargas, C. (2017) Quantitative 3D-imaging for cell biology and ecology of environmental microbial eukaryotes. eLife, 6: e26066.

Daims, H., Lücker, S., Wagner, M. (2006). Daime, a novel image analysis program for microbial ecology and biofilm research. Environ. Microbiol., 8: 200–213.

Haas, LW (1982. Improved epifluorescent microscopic technique for observing planktonic micro-organisms. Ann. Inst. Oceanogr., 58: 261–266.

Hamasaki, K., Long, R. A., and Azam, F. (2004). Individual cell growth rates of marine bacteria, measured by bromodeoxyuridine incorporation. Aquat. Microb. Ecol., 35: 217.

Hatzenpichler, R., Scheller, S., Tavormina, P. L., Babin, B. M., Tirrell, D. A., and Orphan, V. J. (2014). *In situ* visualization of newly synthesized proteins in environmental microbes using amino acid tagging and click chemistry. Environ. Microbiol., 16: 2568–2590.

Hense, B.A., Gais, P., Jutting, U., Scherb, H., Rodenacker, K. (2008) Use of fluorescence information for automated phytoplankton investigation by image analysis. J. Plank. Res., 30: 587–606.

Hobbie, J. E., Daley, R. J., and Jasper, S. (1977). Use of nuclepore filters for counting bacteria by fluorescence microscopy. Appl. Environ. Microbiol., 33, 1225–1228.

Jaqaman, K., Loerke, D., Mettlen, M. et al. (2008). Robust single-particle tracking in live-cell time-lapse sequences. Nat. Methods, 5: 695–702.

Khachikyan, A., Milucka, J., Littmann, S., Ahmerkamp, S., Meador, T., Könneke, M., Burg, T., and Kuypers, M. (2019). Direct Cell Mass Measurements Expand the Role of Small Microorganisms in Nature. Appl. Environ. Microbiol., 85: e00493–19.

Lee, D.W. Hsu, H-L, Bacon, K.E., Daniel, S. (2016). Image restoration and analysis of influenza virions binding to membrane receptors reveal adhesion-strengthening kinetics. PLoS One, 11: e0163437.

Lombard, F., Boss, E., Waite, A.M., et al. (2019). Globally consistent quantitative observations of planktonic ecosystems. Front Mar Sci 6: 196.

McQuin C, Goodman A, Chernyshev V, Kamentsky L, Cimini BA, Karhohs KW, Doan M, Ding L, Rafelski SM, Thirstrup D, Wiegraebe W, Singh S, Becker T, Caicedo JC, and Carpenter AE (2018). CellProfiler 3.0: Next-generation image processing for biology. PLoS Biol., 16: e2005970

Michels, D.E., Lomenick, B., Chou, T-F., Sweredoski, M.J., Pasulka, A.L. (2021). Amino acid analog induces stress response in marine *Synechococcus*. Appl. Environ. Microbiol., 87, e00200–21.

Miloslavich, P., Bax, N.J., Simmons, S.E. et al. (2018). Essential ocean variables for global sustained observations biodiversity and ecosystem changes. Glob. Change Biol., 24: 2416–2433.

Noble, R.T. and Fuhrman, J.A. (1998). Use of SYBR Green I for rapid epifluorescence counts of marine viruses and bacteria. Aquat. Microb. Ecol., 14: 113–118.

Olson, R.J. and Sosik, H.M. (2007). A submersible imaging-in-flow instrument to analyze nano-and microplankton: Imaging FlowCytobot. Limnol. Oceanogr. Methods, 5: 195–203.

Otsu, N. (1979) A threshold selection method from gray-level histograms. IEEE Transactions on Systems, Man, and Cybernetics. 9: 62–66.

Pasulka, A.L., Landry, M.R., Taniguchi, D.A.A., Taylor, A.G., Church, M.J. (2013). Temporal dynamics of phytoplankton and heterotrophic protists at station ALOHA. Deep Sea Res. II, 93: 44–57.

Pasulka, A.L., Thamatrakoln, K., Kopf, S.H., Guan, Y., Poulos, B., Moradian, A., Sweredoski, M.J., Hess, S., Sullivan, M.B., Bidle, K.D., Orphan, V.J. (2018). Interrogating marine virus-host interactions and elemental transfer with BONCAT and nanoSIMS – based methods. Environ. Microbiol., 20: 671–692.

Patel, A., Noble, R.T., Steele, J.A., et al. (2007). Virus and prokaryote enumeration from planktonic aquatic environments by epifluorescence microscopy with SYBR Green I. Nature Protocols, 2: 269–276.

Pernthaler, A., and Amann, R. (2004). Simultaneous fluorescence in situ hybridization of mRNA and rRNA in environmental bacteria. Appl. Environ. Microbiol., 70: 5426–5433.

Polerecky, L., Adam, B., Milucka, J., et al. (2012). Look@NanoSIMS-a tool for the analysis of nanoSIMS data in environmental microbiology. Environ. Microbiol. 14: 1009–1023.

Samo, T.J., Smriga, S., Malfatti, F., Sherwood, B.P., Azam, F. (2014). Broad distribution and high proportion of protein synthesis active marine bacteria revealed by click chemistry at the single cell level. Front Mar. Sci., 1: 48.

Sebastian, M., Gasol, J.M. (2019). Visualization is crucial for understanding microbial processes in the ocean. Phil. Trans. R. Soc. B., 374: 20190083.

Schulze, K., Tillich, U.M., Dandekar, T., Frohme, M. (2013) PlanktoVision—an automated analysis system for the identification of phytoplankton. BMC Bioinformatics, 14: 1.

Sherr, B., and Sherr, E. (1983). Enumeration of heterotrophic microprotozoa by epifluorescence microscopy. Estuarine, Coastal, and Shelf Science, 16: 1–7.

Sherr, B.F., Sherr, E.B., Fallon, R.D. (1987). Use of monodispersed, fluorescently labeled bacteria to estimate in situ protozoan bacterivory. Appl Environ Microbiol., 53:958–965.

Shopov A., Williams, S.C., Verity, P.G. (2000). Improvements in image analysis and fluorescence microscopy to discriminate and enumerate bacteria and viruses in aquatic samples. Aquatic Microbial Ecology, 22: 103–110.

Sosik, H.M. and Olson, R.J. (2007). Automated taxonomic classification of phytoplankton sampled with imaging-in-flow cytometry. Limnol. Oceanogr. Methods, 5: 204–216.

Taylor, A.G., Goericke, R., Landry M.R., Selph, K.E., Wick, D.A., Roadman, M.J. (2012). Sharp gradients in phytoplankton community structure across a frontal zone in the California Current Ecosystem. J Plankton Res., 34: 778–789.

Taylor, A.G., Landry, M.R., Selph, K.E., Wokuluk, J.J. (2015). Temporal and spatial patterns of microbial community biomass and composition in the Southern California Current Ecosystem. Deep Sea Res. II, 112: 117–128.

Turzynski, V., Monsees, I., Moraru, C., Probst, A.J. (2021). Imaging techniques for detecting prokaryotic viruses in environmental samples. Viruses, 13: 2126.

Wait, E.C., Reiche, M.A., Chew, T-L. (2020). Hypothesis-driving quantitative fluorescence microscopy – the importance of reverse-thinking in experimental design. J. Cell Sci., 133: jcp250027.

Wang, I-H., Burckhardt, C.J., Yakimovich, A., Greber, U.F. (2018). Imaging, tracking and computational analysis of virus entry and egress with the cytoskeleton. Viruses, 10, 166.

Waters J. C. (2009). Accuracy and precision in quantitative fluorescence microscopy. The Journal of Cell Biology, 185: 1135–1148.

Waters, J. C., and Wittmann, T. (2014). Concepts in quantitative fluorescence microscopy. Methods in Cell Biology, 123: 1–18.

Weinbauer, M.G., and Suttle, C.A. (1997). Comparison of epifluorescence and transmission electron microscopy for counting viruses in natural marine waters. Aquatic Microbial Ecology, 13, 225–232.

Wollman, R., and Stuurman, N. (2007). High throughput microscopy: from raw images to discoveries. Journal of cell science, 120: 3715–3722.

